# The host antiviral ribonuclease L protein supports Zika virus replication factory formation to enhance infectious virus production

**DOI:** 10.1101/852194

**Authors:** Jillian N Whelan, Joshua Hatterschide, David M. Renner, Beihua Dong, Robert H Silverman, Susan R Weiss

## Abstract

The flavivirus Zika virus (ZIKV) activates ribonuclease L (RNase L) catalytic antiviral function during infection, yet deletion of RNase L decreases ZIKV production, suggesting a proviral role of RNase L. In this study, we reveal that latent RNase L supports ZIKV replication factory (RF) assembly. Deletion of RNase L induced broader cellular distribution of ZIKV dsRNA and NS3 compared with densely concentrated RFs detected in WT cells. An inactive form of RNase L was sufficient to contain ZIKV genome and dsRNA within a smaller area, which increased levels of viral RNA within RFs as well as infectious ZIKV released from the cell. We used a microtubule stabilization drug to demonstrate that RNase L deletion impaired the cytoskeleton rearrangements that are required for proper generation of RFs. During infection with dengue or West Nile Kunjin viruses, RNase L decreased virus production, suggesting that RNase L proviral function is specific to ZIKV.

## Introduction

The flavivirus genus contains arthropod-transmitted viruses with a positive-sense single-stranded RNA (ssRNA) genome, including Zika virus (ZIKV), dengue virus (DENV), and West Nile virus (WNV). These viruses are transmitted by mosquitos and globally distributed, with high associated morbidity and mortality in humans (Wikan and Smith, 2016, Guzman and Harris, 2015, Kaiser and Barrett, 2019). More recently, ZIKV sexual and vertical transmission has been recognized, the latter involving transplacental migration of the virus, potentially resulting in fetal microcephaly (Wu et al., 2016, Cugola et al., 2016, Miner et al., 2016, D’ortenzio et al., 2016, Mlakar et al., 2016, Cauchemez et al., 2016). Due to diversity in ZIKV tissue tropism, disease, and route of transmission as compared with other flaviviruses, it is possible that variances in ZIKV infection at the molecular level confer the observed shifts in clinical outcome at the organismal level. We aimed to identify host machinery that exclusively supports the ZIKV replication cycle, or interactions that are not shared among all flaviviruses, to improve our understanding of the molecular determinants of ZIKV pathogenesis specifically.

After entry into the host cell, the flavivirus genome, which also serves as the mRNA, is directly translated at the ER. Proximal to sites of translation, flaviviruses create replication factories (RFs) through extensive cytoskeletal rearrangements that generate invaginations in the folds of the ER membrane, within which new genome synthesis occurs (Paul and Bartenschlager, 2013, Neufeldt et al., 2018, Cortese et al., 2017, Aktepe et al., 2017, Neufeldt et al., 2019, Welsch et al., 2009). These RFs contain ZIKV replication complex proteins, including the NS3 protease, the replication intermediate dsRNA, as well as template single-stranded genomic RNA. New genome is packaged into compartments at opposite ER folds, and new virions traffic through the trans-golgi network and eventually bud from the plasma membrane (Sager et al., 2018). RFs therefore enable efficient throughput of key viral processes as centers of new genome synthesis linked with viral protein translation as well as new virus assembly. In addition, RFs serve as a protective barrier to impede cytosolic innate immune sensing, as flavivirus RNA predominantly resides within RFs during the bulk of the intracellular replication cycle.

Innate immune sensors within the cytoplasm of the infected cell detect viral RNA to activate antiviral responses, including the type I interferon (IFN) and oligoadenylate synthetase/ribonuclease L (OAS/RNase L) pathways. Extensive research has demonstrated that flaviviruses, including ZIKV, have evolved strategies for counteracting the type I IFN response (Miorin et al., 2017, Riedl et al., 2019, Wu et al., 2017, Bowen et al., 2017, Grant et al., 2016, Best, 2017). Since OAS genes are IFN-stimulated genes (ISGs) and therefore upregulated by type I IFN signaling, the OAS/RNase L pathway can also be potentiated by type I IFN production. However, activation of RNase L can occur in the absence of type I IFN responses when basal OAS expression is sufficient (Whelan et al., 2019, Birdwell et al., 2016). In either event, OAS sensors detect viral dsRNA and generate 2’-5’ oligoadenylates (2-5A). RNase L, which is constitutively expressed in a latent form, homodimerizes upon 2-5A binding to become catalytically active (Dong and Silverman, 1995). Active RNase L cleaves both host and viral ssRNA within the cell (**Figure 1A**, left panel). While there are three OAS isoforms, we have shown that the OAS3 isoform is the predominant activator of RNase L during infection with a variety of viruses including ZIKV (Li et al., 2016, Whelan et al., 2019). Activated RNase L cleavage of host rRNA and mRNA as well as viral ssRNA induces a multitude of events that ultimately inhibits virus infection (Zhou et al., 1997, Castelli et al., 1997, Chakrabarti et al., 2015, Li et al., 2016, Scherbik et al., 2006, Malathi et al., 2007).

**Figure 1.**
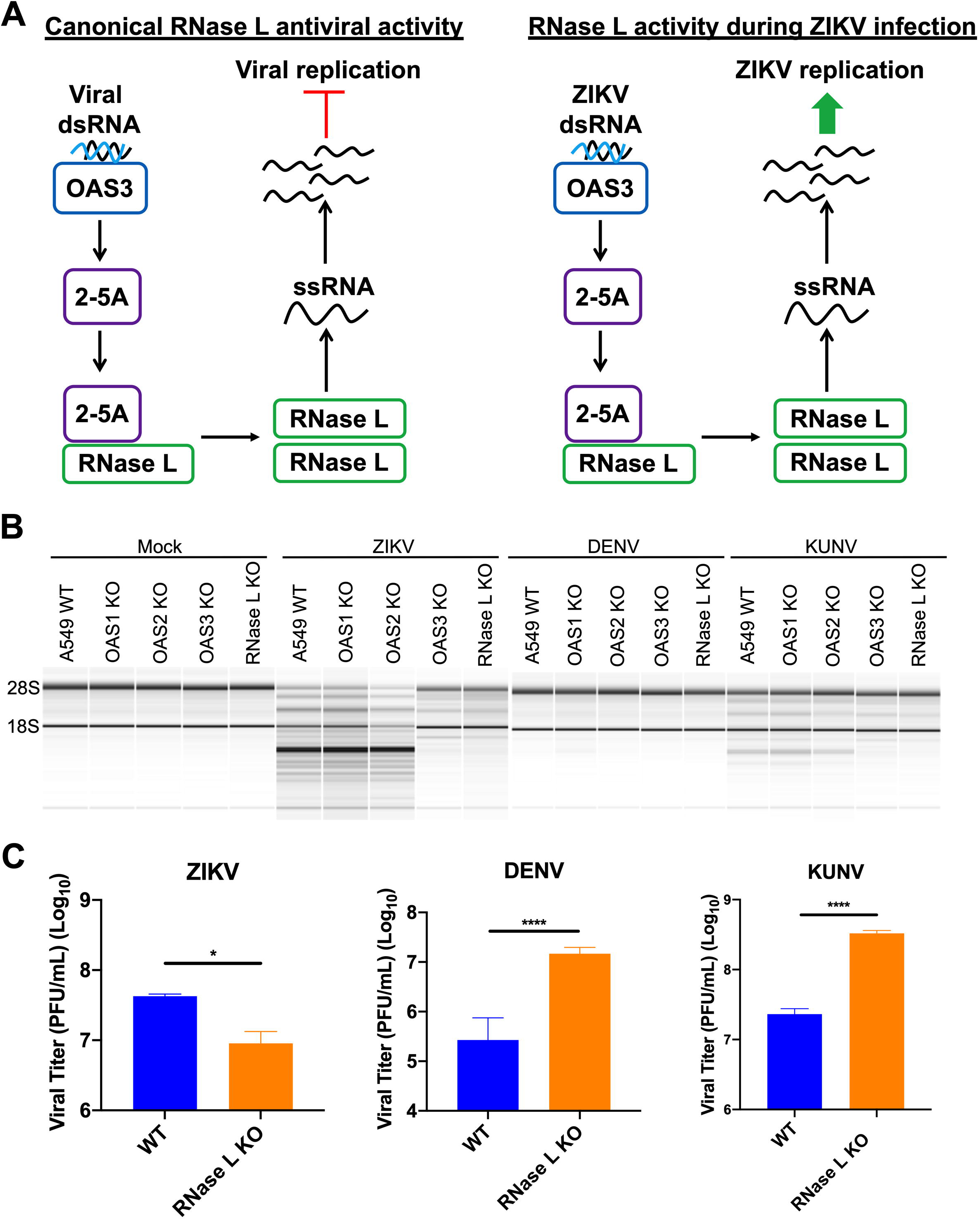
RNase L antiviral activity inhibits DENV and KUNV production but not ZIKV production. (A) Left panel: the canonical RNase L antiviral pathway. Viral dsRNA is detected by OAS3, which produces the small molecule 2-5A that binds latent RNase L, inducing its homodimerization and catalytic activation, resulting in cleavage of host and viral ssRNA, leading to inhibition of viral infection. Right panel: model of RNase L activity during ZIKV infection. ZIKV dsRNA is recognized by OAS3, which activates RNase L resulting in ssRNA cleavage, however ZIKV production is improved with RNase L expression. (B) A549 WT, OAS1 KO, OAS2 KO, OAS3 KO, or RNase L KO cells were mock infected or infected with ZIKV, DENV, or KUNV at an MOI of 1 for 48h. 28S and 18S rRNA integrity was analyzed using an Agilent Bioanalyzer. rRNA degradation is displayed as loss of 28S and 18S rRNA integrity, depicted by a lower banding pattern. (C) A549 WT or RNase L KO cells were infected with ZIKV, DENV, or KUNV at an MOI of 1, supernatants were harvested at 48hpi for measurement of viral titers by plaque assay, shown as plaque forming units (PFU)/mL virus. Data is representative of at least two independent experiments. Statistical significance was determined by Student’s t test, displayed is the mean of three replicates ± SD, *p<0.05, ****p<0.0001.

Once activated, RNase L can restrict infection of a diverse range of DNA and RNA viruses, including flaviviruses DENV and WNV (Silverman, 2007, Lin et al., 2009, Scherbik et al., 2006, Samuel et al., 2006). Many viruses have subsequently developed mechanisms for evading RNase L antiviral effects, most of which target this pathway upstream of RNase L activation through degradation or sequestration of OAS proteins or degradation of 2-5A (Silverman, 2007, Zhao et al., 2012, Drappier et al., 2018, Silverman and Weiss, 2014, Sanchez-Tacuba et al., 2015, Goldstein et al., 2017, Zhang et al., 2013). We recently demonstrated that ZIKV avoids antiviral effects of activated RNase L, and that this evasion strategy required assembly of RFs to protect genome from RNase L cleavage (Whelan et al., 2019). Despite substantial RNase L-mediated cleavage of intracellular ZIKV genome, a portion of uncleaved genome was shielded from activated RNase L within RFs. This genome was sufficient to produce high levels of infectious virus particles, as infectious ZIKV released from WT cells was significantly higher than from RNase L KO cells. These results indicated that RNase L expression was ultimately proviral during ZIKV infection (**Figure 1A**, right panel). Unlike ZIKV, DENV RFs did not obstruct antiviral RNase L activity, and infectious virus production was instead decreased in the presence of RNase L (Whelan et al., 2019, Lin et al., 2009). As this was the initial report of viral resistance to catalytically active RNase L during infection, we sought to isolate the differences between ZIKV RFs and those constructed by other flaviviruses, to identify factors that enable this ZIKV evasion mechanism.

In this study, we focused on elucidating how RNase L increases ZIKV production. An earlier study reported latent RNase L operating as a component of the actin cytoskeleton to reorganize cellular framework during viral infection (Malathi et al., 2014). Since it is well understood that flaviviruses reorganize the cellular cytoskeletal and organellar network during infection (Neufeldt et al., 2018), we investigated the possibility that RNase L association with the cytoskeleton was exploited by ZIKV to assemble highly protective RFs that dually served as a barrier against host sensors in addition to providing sites of replication.

## Results

### RNase L antiviral activity inhibits DENV and WNV production but not ZIKV production

We have previously shown that ZIKV activation of the OAS/RNase L pathway can be detected by 24hpi (Whelan et al., 2019). Using host 28S and 18S rRNA degradation as a measurement for RNase L activation, we show that RNase L activation during ZIKV infection is mediated by the OAS3 isoform specifically (**Figure 1A**, left panel). By 48hpi, degradation of rRNA was detected in ZIKV-infected OAS1 KO and OAS2 KO cells similar to that of parental WT cells, while KO of OAS3 or RNase L prevented as well as confirmed RNase L-mediated cleavage of rRNA (**Figure 1B**). In contrast to other RNase L-activating viruses, RNase L nuclease activity did not restrict ZIKV production. This was demonstrated by a lack viral titer increase by 48hpi, after multiple rounds of replication, in ZIKV-infected RNase L KO cells compared to in WT cells (**Figure 1C**). Instead, deletion of RNase L reduced ZIKV production. In comparison to ZIKV, we detected weaker activation of RNase L by West Nile virus Kunjin strain (KUNV), although also OAS3-dependent, and could not detect any RNase L activation by DENV using this assay (**Figure 1B**). Despite reduced RNase L activation by DENV and KUNV, both viruses were restricted by RNase L activity in WT cells (**Figure 1C**). Therefore, RNase L is not only ineffective in inhibiting ZIKV infection, but also specifically promotes ZIKV production (**Figure 1A**, right panel).

### RNase L improves ZIKV RF function to increase ZIKV RNA and protein expression

Since infectious virus production was enhanced by RNase L expression, we examined whether RNase L was required for optimal ZIKV replication. We used immunofluorescence assays (IFAs) for detection of the replication intermediate dsRNA and the ZIKV NS3 protein as markers for ZIKV RFs, comparing expression levels in WT and RNase L KO cells. At 20hpi, expression of both ZIKV dsRNA and NS3 was increased in WT cells, as measured by mean dsRNA and NS3 intensity at perinuclear ER sites characteristic of flaviviruses (**Figure 2AB&C**). We also observed a change in localization of ZIKV dsRNA and NS3 in RNase L KO cells, both of which were more disseminated around the nucleus and into the cytoplasm. We used circularity and diameter parameters to quantify the spread of viral products throughout the cell at 20hpi, and determined that RNase L deletion decreased circularity and increased diameter of RFs, distinct from the densely concentrated RF phenotype observed in WT cells (**Figure 2AB&C**). We observed this trend of viral RF diffusion in RNase L KO cells as early as 16hpi through 30hpi, in two different RNase L KO clones (**Figure S1**). We performed all future analyses on cells fixed at 20hpi, which provides adequate infected cells for quantification purposes, but is early enough for examination of a uniform stage of the viral replication cycle across each sample. Additionally, since alterations in ZIKV dsRNA and NS3 were nearly identical, we proceeded to measure effects on RFs using dsRNA expression only.

**Figure 2.**
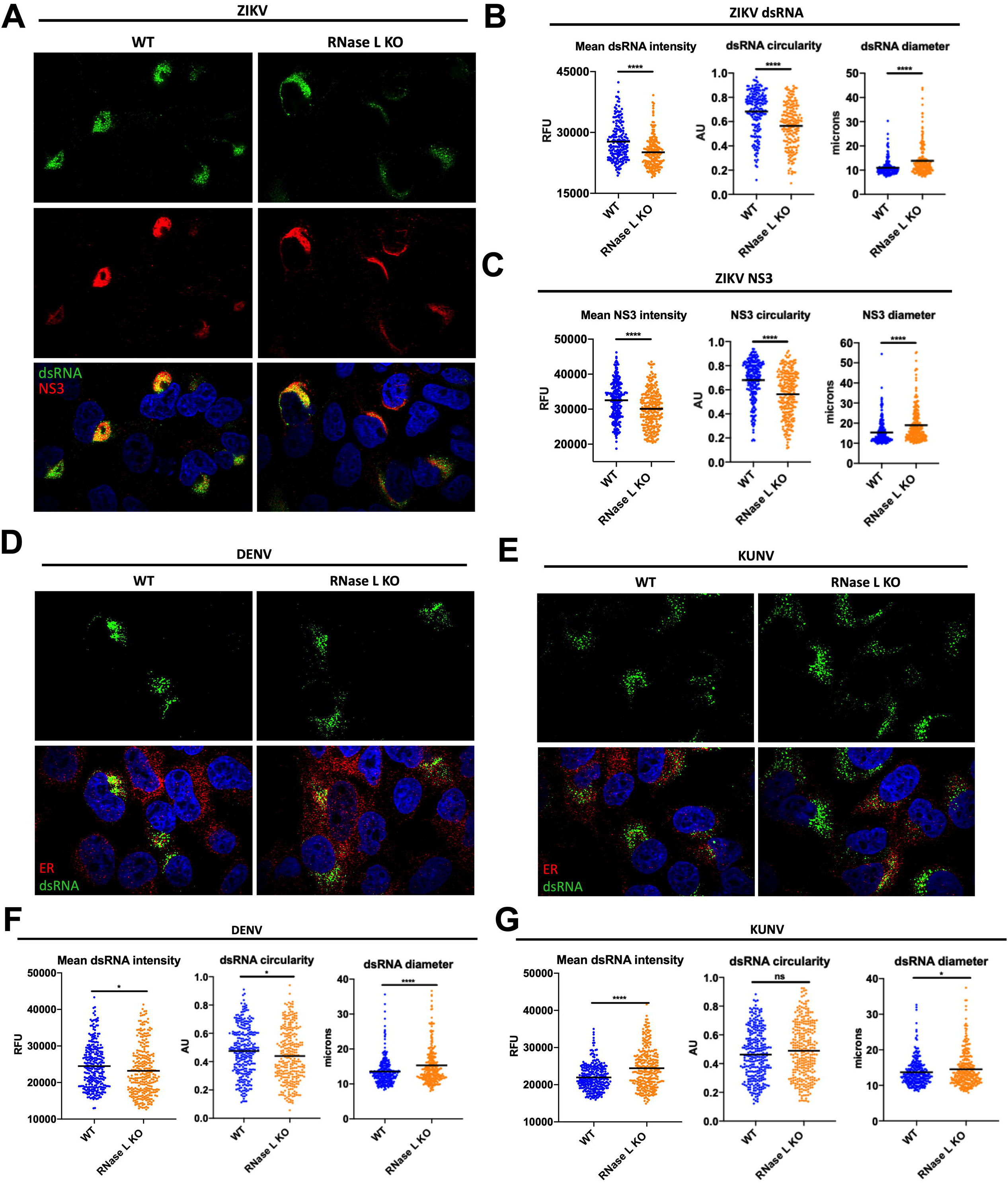
RNase L improves RF function to increase ZIKV RNA and protein expression. A549 WT and RNase L cells were infected at an MOI of 1, cells were fixed at 20hpi for immunofluorescence assays (IFA). (A) ZIKV infected cells were stained for dsRNA (green) and ZIKV NS3 (red), with DAPI (blue) staining of nuclei. Quantification of ZIKV dsRNA (B) and NS3 (C) mean intensity, circularity, and diameter of staining shown in (A). (D) DENV or (E) KUNV infected cells were stained for dsRNA (green) and ER (PDI, red), with DAPI (blue) staining of nuclei. Quantification of (F) DENV or (G) KUNV dsRNA mean intensity, circularity, and diameter of staining shown in (D) and (E), respectively. Data is representative of at least two independent experiments. Statistical significance was determined by Student’s t test, black bars represent the mean, ns = not significant, *p<0.05, ****p<0.0001. Imaged at 100X magnification. RFU = relative fluorescence units, AU = arbitrary units. See also Figure S1.

When comparing DENV dsRNA expression and localization in RNase L KO cells to that of WT cells, we found that dsRNA intensity and circularity were decreased while diameter was increased without RNase L, a similar trend but to a lesser degree than with ZIKV (**Figure 2D&F**). Furthermore, this does not result in a defect in virus release as measured by viral titer, but instead virus titer is enhanced by RNase L KO (**Figure 1C**), suggesting slight changes in RF intensity and shape at the ER do not translate to changes in virus titer during DENV infection. In contrast, KUNV dsRNA intensity increased in RNase L KO cells compared to WT, with little or no change in diameter or circularity of dsRNA, respectively (**Figure 2E&G**).

### Effects of OAS3 knockout on ZIKV RF function

To determine if the involvement of RNase L in ZIKV RF structure requires its nuclease function, we first evaluated ZIKV dsRNA levels and shape of RFs in OAS3 KO cells. As shown in Figure 1, ZIKV activation of RNase L is dependent on OAS3 expression, therefore deletion of OAS3 eliminates RNase L catalytic activity without affecting RNase L levels (Li et al., 2016). To confirm RNase L latency in OAS3 KO cells, we measured 2-5A, the activator of RNase L synthesized by OAS enzymes, with a fluorescence resonance energy transfer (FRET) assay during ZIKV infection of WT, RNase L KO, and OAS3 KO cells. We detected 2-5A generation during ZIKV infection of both WT and RNase L KO cells, however 2-5A in ZIKV-infected OAS3 KO cells was not increased over levels in mock infected WT cells (**Figure 3A**). This confirmed previous results indicating that OAS3 is the primary human OAS isoform required for 2-5A synthesis during viral infections (Li et al., 2016, Whelan et al., 2019), and verified that other OAS isoforms did not compensate for loss of OAS3 expression during ZIKV infections. Together these results suggest that OAS3 is the primary producer of 2-5A during ZIKV infection, thereby providing a useful system for investigating RNase L function independent of its catalytic activity during viral infection.

**Figure 3.**
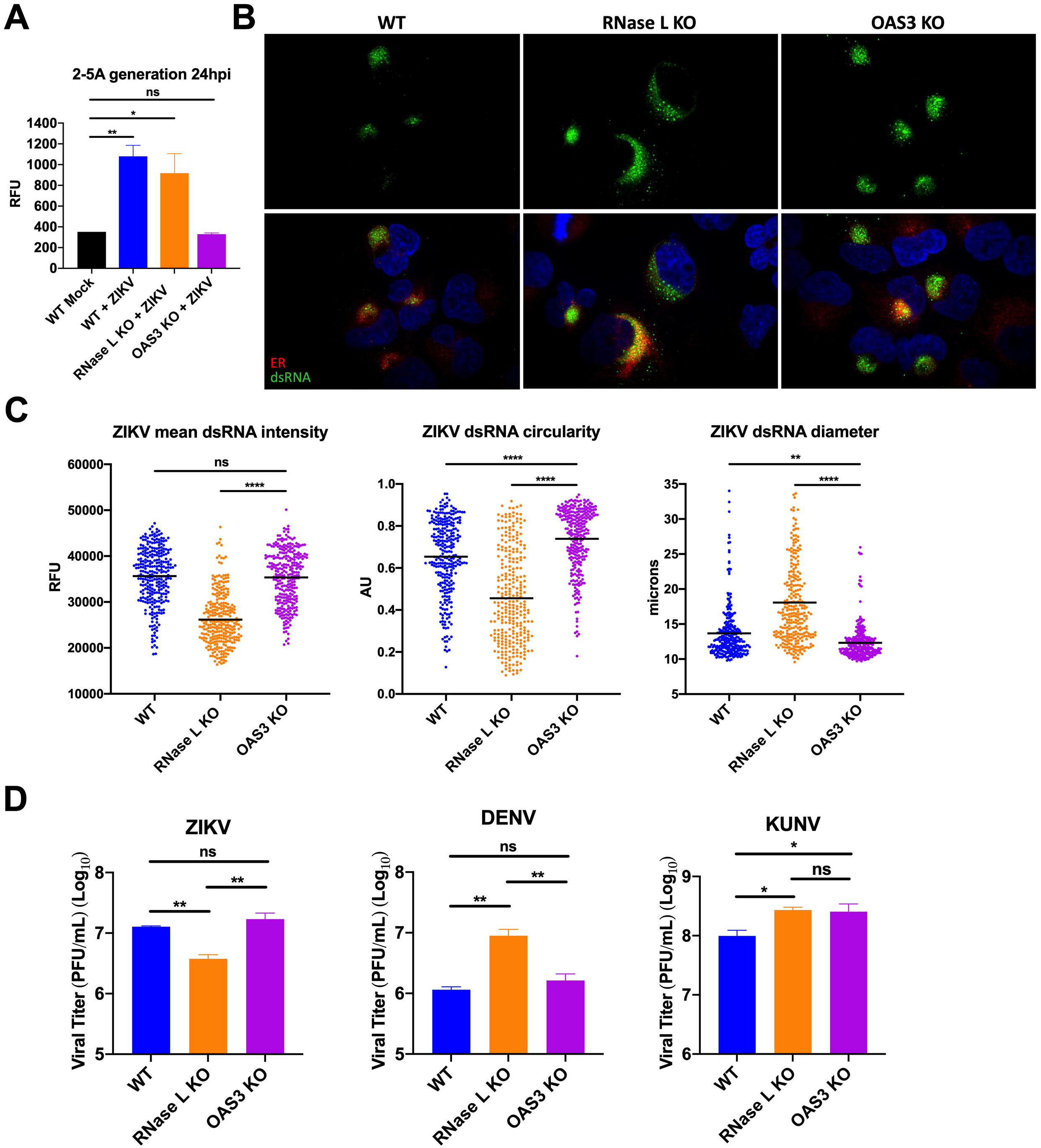
Effects of OAS3 knockout on ZIKV RF function. (A) A549 WT, RNase L KO, or OAS3 KO cells were mock infected or infected with ZIKV at an MOI of 5, 24hpi lysates were harvested for 2-5A FRET assay. (B) A549 WT, RNase L KO, or OAS3 KO cells were infected with ZIKV at an MOI of 1, fixed at 20hpi, and stained for dsRNA (green) and ER (PDI, red), with DAPI (blue) staining of nuclei. (C) Quantification of ZIKV mean intensity, circularity, and diameter of staining shown in (B). (D) A549 WT, RNase L KO, or OAS3 KO cells were infected with ZIKV, DENV, or KUNV at an MOI of 0.1, supernatants were harvested at 48hpi for measurement of viral titers by plaque assay, shown as plaque forming units (PFU)/mL virus. Data is representative of at least two independent experiments. Statistical significance was determined by one-way ANOVA, displayed is the mean of three replicates ± SD for replication assays, ns = not significant, *p<0.05, **p<0.01, ****p<0.0001. Imaged at 100X magnification. RFU = relative fluorescence units, AU = arbitrary units. See also Figure S2.

We next used IFAs to compare dsRNA expression levels and localization in ZIKV infected OAS3 KO cells to that of infected WT and RNase L KO cells. We used protein disulfide isomerase (PDI) as a marker for the ER, where RFs reside. At 20hpi, ZIKV dsRNA intensity in the OAS3 KO cells was rescued to WT levels from that of RNase L KO cells (**Figure 3B&C**). Furthermore, RFs in OAS3 KO cells were more circular and smaller in diameter than in both RNase L KO and WT cells, suggesting additional RF support from inactive RNase L compared to in WT cells (**Figure 3B&C**). While some changes in DENV and KUNV dsRNA expression in OAS3 KO cells displayed a similar trend as that of ZIKV dsRNA, changes were minimal in comparison with alterations in ZIKV dsRNA observed in OAS3 KO cells (**Figure S2**).

We also found that by 48hpi OAS3 KO, which suppresses RNase L activation, restored infectious ZIKV production from reduced levels of RNase L KO cells to that observed during WT cell infection (**Figure 3D**). DENV production, which was limited by RNase L in WT cells, was not improved by OAS3 KO, suggesting an OAS3-independent RNase L-mediated inhibition of DENV infection (**Figure 3D**). While KUNV production was also limited by RNase L in WT cells, it improved with OAS3 KO as observed with ZIKV. However, unlike DENV, KUNV activation of RNase L was OAS3-dependent (Figure 1B). Therefore, increased KUNV yields from OAS3 KO cells were likely due to the lack of RNase L mediated antiviral activity (**Figure 3D**). These results from cells lacking OAS3 suggest that ZIKV infection may utilize an inactive form of RNase L in a unique manner that results in higher virus production than with RNase L absent.

### RNase L maintains ZIKV RF structure in the absence of OAS3 to enhance virus replication

Viral dsRNA expression and localization informs on replication levels and RF location in the cell, respectively, but does not entirely reflect changes in viral genome. While dsRNA activates the OAS/RNase L pathway, ssRNA including viral genome is the target of RNase L nucleolytic activity. Using fluorescence *in situ* hybridization (FISH) to visualize ZIKV genome, we detected an RNase L-dependent absence of ZIKV genome outside of RFs as early as 20hpi, which we attributed to RNase L cleavage of genome (Whelan et al., 2019). To understand RNase L non-catalytic effects on viral genome, we examined ZIKV genome in OAS3 KO cells, using PDI staining as a marker for RF sites. At 20hpi, ZIKV genomic RNA in WT cells was only detectable at or near the ER. Genome in RNase L KO cells was at the ER, but also dispersed throughout the cell. In OAS3 KO cells, genome remained at or proximal to the ER as observed in WT cells, despite the apparent absence of activated RNase L (**Figure 4A**). While some genome was detected outside of the ER in OAS3 KO cells, this extra-RF genome was minimal compared to that of RNase L KO cells, and possibly genome otherwise cleaved by activated RNase L in WT cells (**Figure 4A**). Quantification of total genome staining within infected cells revealed that deletion of RNase L did not alter intensity of genome expression when compared to WT cells. Since genome staining was not circular in shape, we instead used total area to measure increased dissemination of genome throughout the infected cell without RNase L (**Figure 4D**). Inactive RNase L in OAS3 KO cells restored the area of genome expression from that of RNase L KO cells back to the area measured in WT cells (**Figure 4D**). Furthermore, genome confined to RFs in OAS3 KO cells resulted in higher levels of genome compared to WT and RNase L KO cells (**Figure 4D**), suggesting that genome location within the cell dictates new virus release (Figure 3D). Enhanced genome intensity in OAS3 KO cells over that of WT cells is possibly due to inactive RNase L availability for RF support, as opposed to in WT cells wherein RNase L is activated and performs other functions, namely degrading RNA.

**Figure 4.**
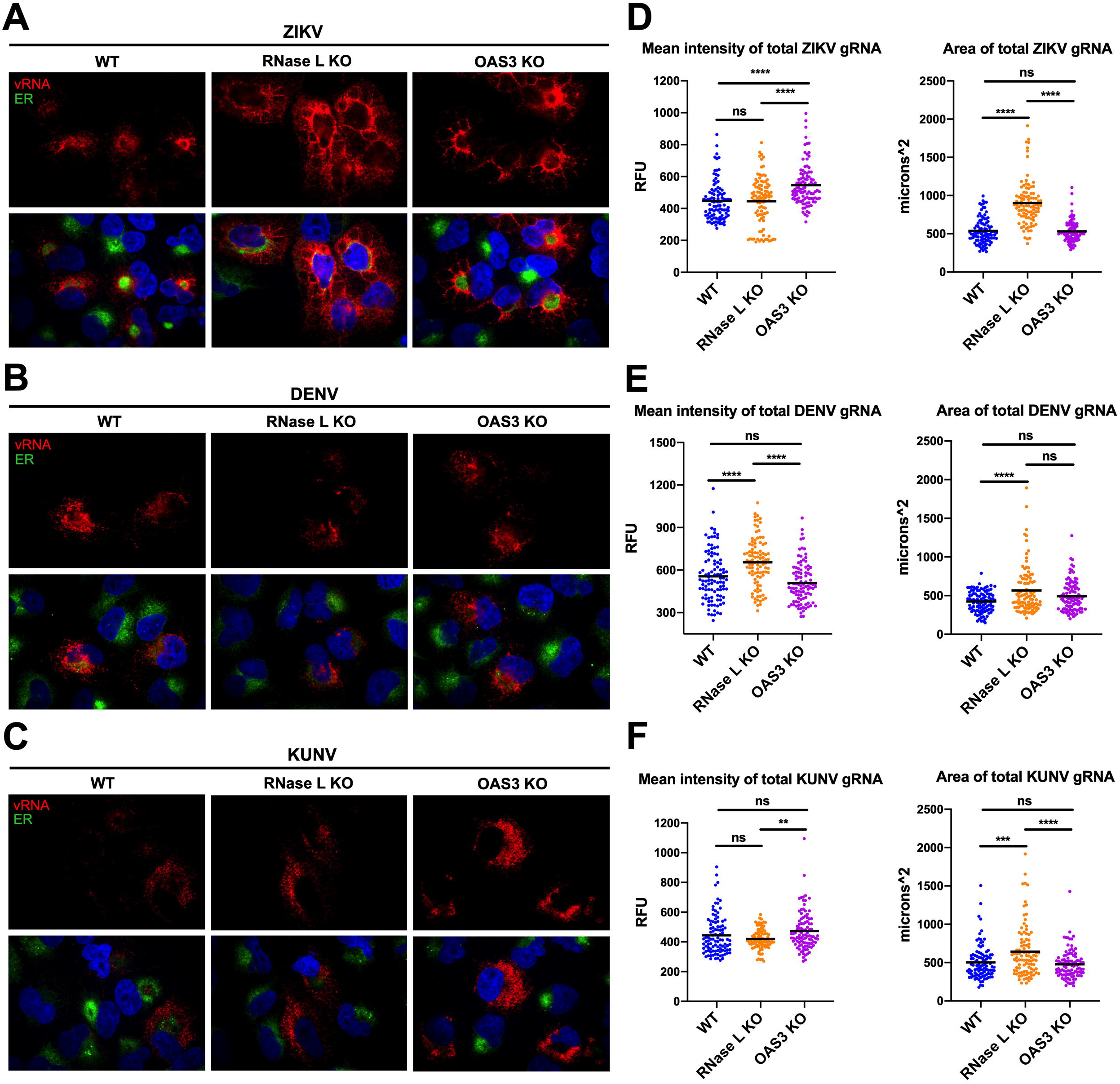
RNase L maintains ZIKV RF structure in the absence of OAS3 to enhance virus replication. A549 WT, RNase L KO, or OAS3 KO cells were infected at an MOI of 1 with (A) ZIKV, (B) DENV, or (C) KUNV. Cells were fixed at 20hpi and stained with a virus genome-specific probe (red) using fluorescence *in situ* hybridization, before staining ER (PDI, green) and nuclei with DAPI (blue) by IFA. Quantification of the mean intensity and area of (D) ZIKV, (E) DENV, or (F) KUNV genome staining. Data is representative of at least two independent experiments. Statistical significance was determined by one-way ANOVA, ns = not significant, **p<0.01, ***p<0.001, ****p<0.0001. Imaged at 100X magnification. RFU = relative fluorescence units.

In contrast to ZIKV, DENV and KUNV genome staining and cellular localization was similar in WT, RNase L KO, and OAS3 KO cells, with genome residing at the ER regardless of RNase L presence or activation status (**Figure 4B&C**). Quantification of DENV total genome staining determined that RNase L deletion increased the area and intensity of genome expression over that in cells lacking OAS3-dependent RNase L activity (**Figure 4E**), suggesting OAS3-independent antiviral effects of RNase L on DENV new genome synthesis, which correlates with increased DENV titers in RNase L KO cells but not OAS3 KO cells (Figure 3D). Quantification of KUNV genome staining exhibited only minor alterations in intensity, with genome in OAS3 KO cells only increased over that in RNase L KO cells, but not increased over that in WT cells as was observed for ZIKV (**Figure 4F**). Since KUNV does not activate RNase L until 48hpi by rRNA degradation assay (Figure 1B), we had not expected a change in genome intensity between cell lines at 20hpi. Additionally, the area of KUNV genome expression was increased in RNase L KO cells compared to both WT and OAS3 KO cells, but to a lesser degree than during ZIKV infection (**Figure 4F**). Overall, absence of OAS3-dependent RNase L activity had a greater impact on ZIKV genome localization and expression compared to that of DENV and KUNV, which agrees with its minimal effects on DENV and KUNV RFs and viral titers.

### ZIKV RFs are unaffected by pIC treatment in the absence of OAS3

The lack of rRNA degradation (Figure 1B) and 2-5A production (Figure 3A) during ZIKV infection of OAS3 KO cells indicated that OAS1 and OAS2 did not compensate for OAS3 in its absence. Despite this evidence, we wanted to further investigate whether ZIKV genome localization during infection of OAS3 KO cells (Figure 4A&D) was due to non-catalytic, proviral effects of RNase L on RF structure, and not a result of activated RNase L cleavage of ZIKV genome residing outside of RFs. As an alternative assay for RNase L activation during ZIKV infection, we used the synthetic dsRNA poly(rI):poly(rC) [pIC] to artificially activate RNase L prior to ZIKV infection, which induces RNase L-mediated cleavage of viral genome before RFs are assembled during infection. Thus, we can use dsRNA expression as a readout for RF assembly, which would be reduced after pIC treatment in cells where RNase L is active. We examined RF presence at 24hpi when RNase L would be activated by ZIKV infection alone (Whelan et al., 2019). RFs were apparent in cells infected with ZIKV without any pIC treatment, demonstrated by the presence of dsRNA IFA staining in WT cells transfected with lipofectamine only (**Figure 5A**, left panel). Transfection of pIC 2h before ZIKV infection robustly activates multiple innate immune pathways to block virus replication, including RNase L. In WT cells, pIC treatment preceding ZIKV infection resulted in the lack of detectable dsRNA (**Figure 5A**, right panel). In RNase L KO cells, dsRNA was detectable, validating that absence of dsRNA in WT cells was due to RNase L specifically and not from other antiviral responses induced by pIC (**Figure 5B**). In OAS3 KO cells, pIC transfection before ZIKV infection had no effect on dsRNA expression, further demonstrating that RNase L was not activated in the absence of OAS3 (**Figure 5C**).

**Figure 5.**
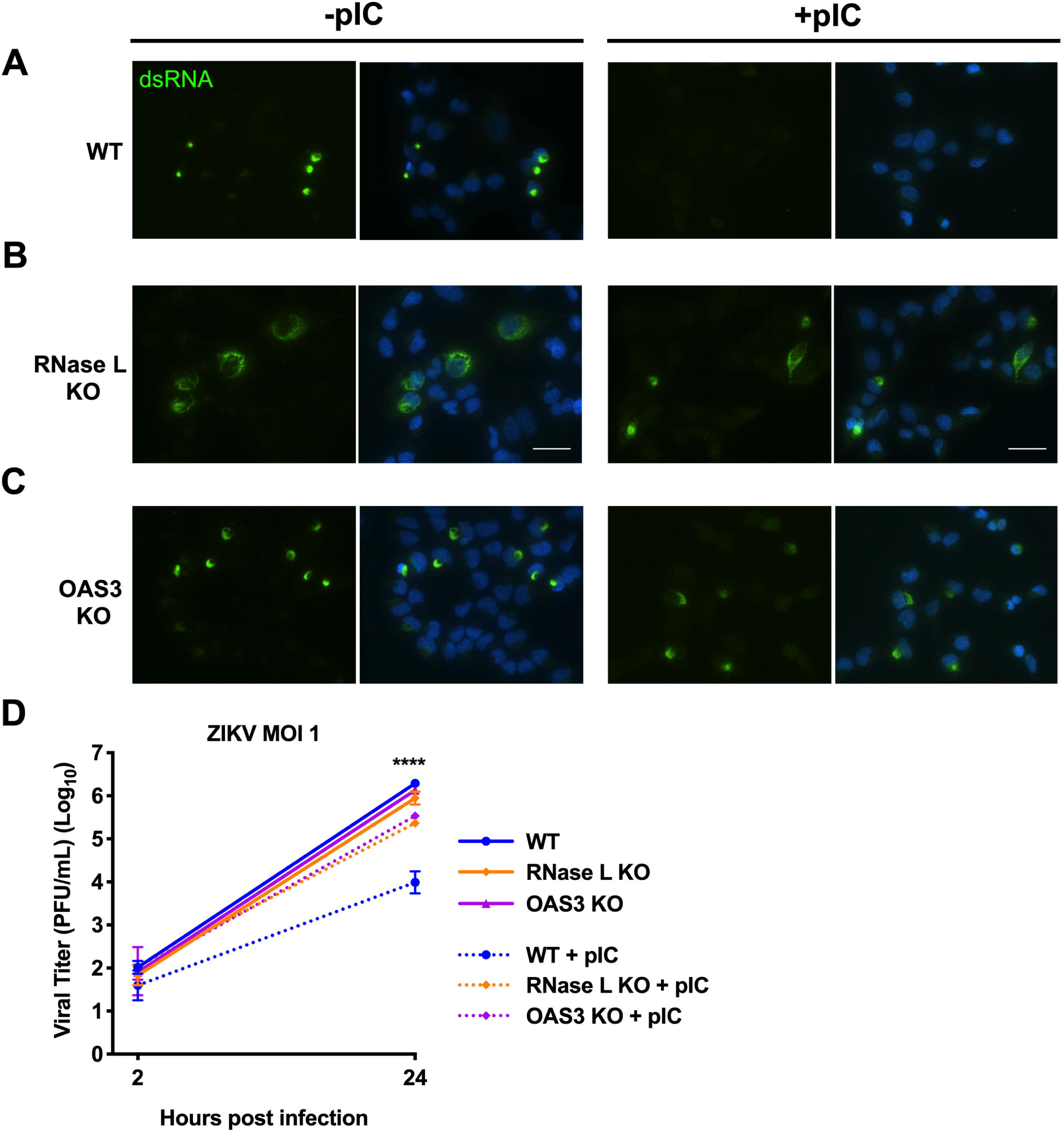
ZIKV RFs are unaffected by pIC treatment in the absence of OAS3. A549 WT, RNase L KO, or OAS3 KO cells were transfected with lipofectamine only or with pIC for 2h before infection with ZIKV at an MOI of 1. (A) Infected cells were fixed at 24hpi, and stained by IFA for dsRNA (green), with DAPI for nuclei staining (blue). (B) Supernatants from infected cells was harvested at 2 or 24hpi for measurement of viral titers over time by plaque assay, shown as plaque forming units (PFU)/mL virus. Data is representative of at least two independent experiments. Statistical significance was determined by one-way ANOVA, displayed is comparison of WT ± pIC (****), and the mean of three replicates ± SD, ****p<0.0001. Imaged at 60X magnification, scale bar 500μm.

To corroborate these results with a more quantitative method, we also used this system to demonstrate that pIC treatment before ZIKV infection of OAS3 KO cells had no effect on infectious ZIKV production. After transfecting cells for 2h with pIC, cells were infected with ZIKV for 24hpi, and viral titers were measured by plaque assay. Without pIC, ZIKV titers in all three cell lines were similar at 24hpi, which agrees with previous data showing decreased ZIKV from RNase L KO cells only after 24hpi (**Figure 5D**). In WT cells with pIC-induced activation of RNase L, infectious ZIKV production was restricted. In RNase L KO and OAS3 KO cells, pIC only slightly inhibited ZIKV production, indicating that restriction of viral titers in WT cells was predominantly due to RNase L antiviral activity (**Figure 5D**). These results correlate with absence of dsRNA detection in only WT cells after pIC transfection (**Figure 5C**), and establish that RNase L is not catalytically active in OAS3 KO cells. This also maintains that decreased ZIKV in RNase L KO cells compared to WT and OAS3 KO cells (Figure 3D) is not due to RNase L-independent antiviral responses that subsequently limit virus production, as responses unrelated to RNase L induced by pIC in RNase L KO or OAS3 KO cells had only a minor effect on titers (**Figure 5D**). Finally, these results suggest that inactive RNase L retains ZIKV genome in smaller, more efficient RFs.

### Expression of RNase L catalytic mutant R667A in HeLa M cells enhances ZIKV RF function and virus production

We next directly investigated whether an inactive form of RNase L can enhance ZIKV RF function to augment infectious virus production using a different cell line. To complement our A549 model, in which RNase L pathway components were deleted, we infected HeLa M cells stably expressing either RNase L WT (+RL WT) or RNase L nuclease dead mutant R667A (+RL R667A) (Dong et al., 2001). We also infected HeLa M cells expressing the empty pcDNA3 vector (vector control, VC) that have undetectable levels of RNase L protein by western blotting (**Figure 6A**). This provided an RNase L-deficient cell line as a control for this system, in addition to verification that effects in RL R667A-expressing cells were not possibly due to endogenous RNase L WT expression. While RL WT protein expression was lower than that of RL R667A, it was sufficient for RNase L-mediated rRNA degradation, which was only detectable in ZIKV infected +RL WT cells and not VC or +RL R667A cells, despite OAS3 protein expression in all three cell lines (**Figure 6A&B**). We evaluated effects of RNase L WT or R667A expression on ZIKV dsRNA expression and localization compared to that of VC cells at 20hpi, using PDI ER staining to denote RF sites. We found that expression of either RL WT or RL R667A increased ZIKV dsRNA intensity at the ER over that in VC cells, however only RL R667A increased circularity of ZIKV RFs in comparison to those of VC cells (**Figure 6C&D**). To determine if RNase L effects on ZIKV dsRNA intensity or circularity correlated with elevated infectious ZIKV titers, we measured virus from infected HeLa M cells at 48hpi and detected a significant increase in ZIKV production in RL R667A-expressing cells compared to VC or RL WT-expressing cells (**Figure 6E**). In addition, we found that neither RL WT nor RL R667A expression increased DENV or KUNV titers, demonstrating a proviral effect of catalytically inactive RNase L on ZIKV in particular (**Figure 6E**). Furthermore, expression of RL WT, but not RL R667A, slightly decreased KUNV production at 48hpi (**Figure 6E**). As RNase L is activated during KUNV infection at this timepoint, restricted virus production is likely a result of RNase L catalytic activity to inhibit KUNV replication.

**Figure 6.**
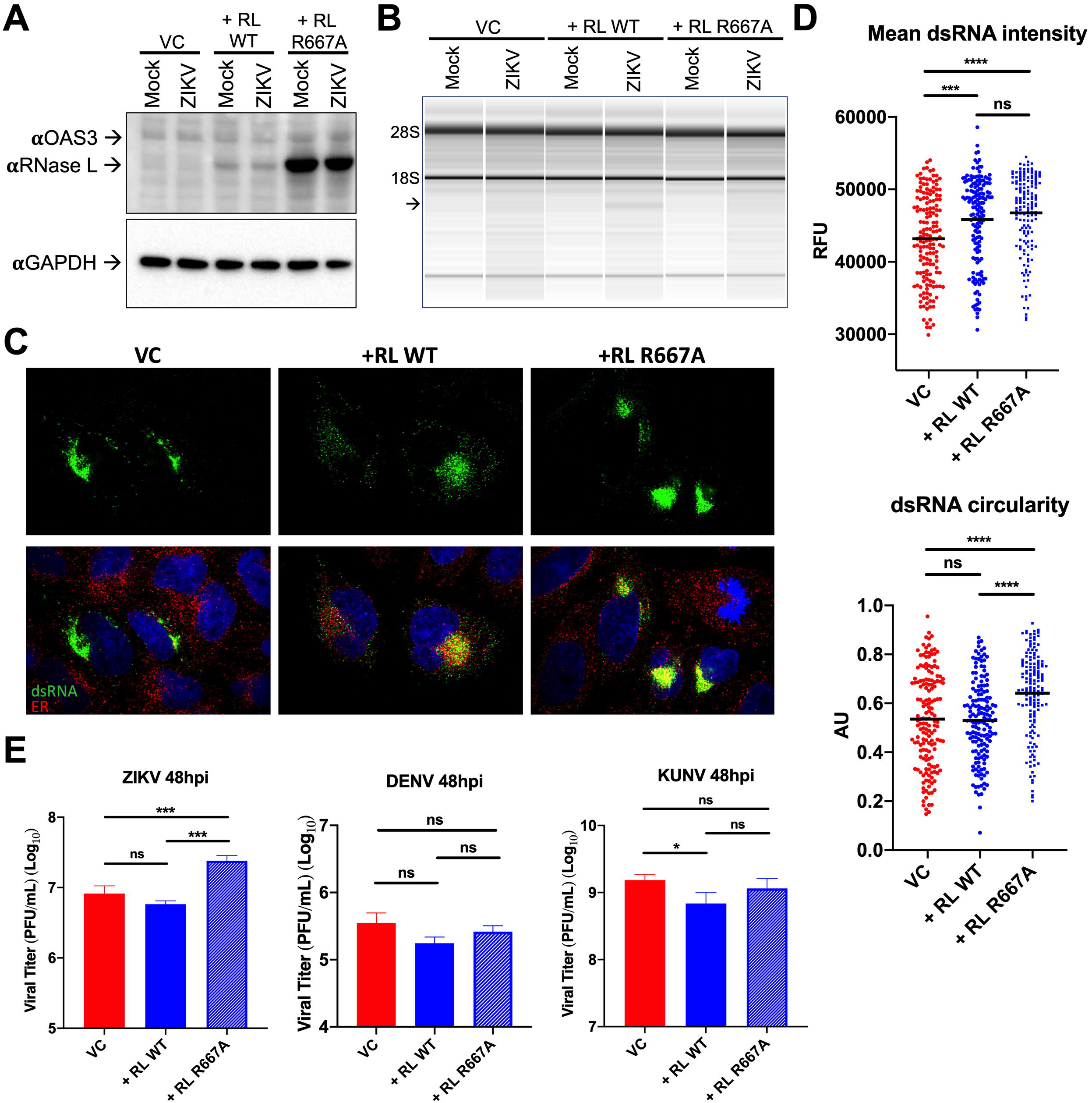
Expression of RNase L catalytic mutant R667A in HeLa M cells enhances ZIKV RF function and virus production. HeLa M cells expressing empty vector (vector control, VC), RNase L WT (RL WT), or RNase L nuclease dead mutant R667A (RL R667A) were mock infected or infected with ZIKV at and MOI of 10. Protein lysates were harvested at 24hpi and OAS3, RNase L, and GAPDH (loading control) were detected by western blotting. (B) HeLa M cells were mock infected or infected with ZIKV at an MOI of 5. RNA was harvested 48hpi, and 28S and 18S rRNA integrity was analyzed using an Agilent Bioanalyzer. rRNA degradation is displayed as loss of 28S and 18S rRNA integrity, depicted by a lower banding pattern (black arrow). (C) HeLa M cells were infected with ZIKV at an MOI of 1. At 20hpi, cells were fixed and stained for dsRNA (green) and ER (PDI, red), with DAPI (blue) staining of nuclei. (D) Quantification of ZIKV mean intensity and circularity shown in (C). (E) HeLa M cells were infected with ZIKV, DENV, or KUNV at an MOI of 0.1, supernatants were harvested at 48hpi for measurement of viral titers by plaque assay, shown as plaque forming units (PFU)/mL virus. Data is representative of at least two independent experiments. Statistical significance was determined by one-way ANOVA, displayed is the mean of three replicates ± SD for replication assays, ns = not significant, *p<0.05, ***p<0.001, ****p<0.0001. Imaged at 100X magnification. RFU = relative fluorescence units, AU = arbitrary units.

### Effects of RNase L deletion on ZIKV RFs resemble antiviral effects of microtubule stabilization on ZIKV RFs

There have been multiple reports of RNase L interactions with the host cytoskeleton, some of which are contingent upon RNase L catalytic latency (Malathi et al., 2014, Tnani et al., 1998). Since ZIKV rearrangement of the host cytoskeleton to form RFs within ER invaginations is crucial to ZIKV infection, we hypothesized that inactive RNase L association with the cytoskeleton is exploited by ZIKV to form RFs. Without an effective antibody for IFA detection of RNase L, we were unable to visualize RNase L localization during ZIKV infection, or if this localization changed upon catalytic activation. Instead, to demonstrate the role of RNase L in ZIKV RF formation, we treated ZIKV infected cells with paclitaxel, a microtubule stabilizing drug, and compared its effects on ZIKV RFs with effects of RNase L deletion. Paclitaxel has been shown to inhibit ZIKV titers (Cortese et al., 2017), likely by blocking ZIKV-induced microtubule rearrangements for establishing RFs, although the mechanism by which paclitaxel inhibits ZIKV infection has not been shown. Microtubule stabilization mediated by paclitaxel would prohibit rearrangements necessary for forming ZIKV RFs. We predicted that RNase L deletion similarly impaired ZIKV-stimulated cytoskeletal rearrangements for RF assembly, thereby reducing virus production.

A549 cells were treated with paclitaxel or vehicle (DMSO) 3h after infection with ZIKV. Cells were fixed at 20hpi and stained for ZIKV dsRNA and NS3 by IFA. Expression of ZIKV dsRNA and NS3 in DMSO-treated RNase L KO cells was again dimmer and more dispersed throughout the cell compared to in WT cells (**Figure 7A**). Quantification confirmed that the absence of RNase L lowered ZIKV dsRNA expression and concentration, as measured by intensity and circularity of dsRNA staining, respectively (**Figure 7C**). In paclitaxel treated WT cells, ZIKV dsRNA and NS3 staining resembled that of RNase L KO with or without paclitaxel (**Figure 7B**). Quantification maintained a decrease in dsRNA intensity and circularity in WT cells treated with paclitaxel, closer to levels of RNase L KO cells, compared to DMSO-treated WT cells. Additionally, paclitaxel had no effect on dsRNA intensity and minimal impact on circularity in RNase L KO cells (**Figure 7C**). We validated effects of paclitaxel treatment on microtubule stability by staining βtubulin, which together with αtubulin makes up microtubules. In ZIKV infected WT cells, βtubulin localized to dsRNA staining, which agrees with previous findings of ZIKV-induced microtubule rearrangements surrounding RFs (**Figure 7D**) (Cortese et al., 2017). In RNase L KO cells, βtubulin was no longer detectable at concentrated RFs, similar to dissemination of ZIKV dsRNA or NS3 without RNase L expressed (**Figure 7D**). Paclitaxel treatment of both ZIKV infected WT and RNase L KO cells resulted in βtubulin aggregation due to inhibition of microtubule tractability (**Figure 7E**). Finally, we treated cells with paclitaxel 3h after infecting with ZIKV to measure changes in infectious virus production at 24hpi as a result of microtubule stabilization in WT and RNase L KO cells. Paclitaxel in WT cells reduced infectious ZIKV titers to the level of RNase L KO cells (**Figure 7F**). Moreover, paclitaxel did not reduce ZIKV titers in RNase L KO cells, suggesting that the paclitaxel means of blocking ZIKV production was lost in RNase L KO cells where microtubule associations with ZIKV RFs were already weak due to absence of RNase L (**Figure 7F**). Together with the effects of paclitaxel on dsRNA expression, these results offer a mechanism for RNase L-mediated increased ZIKV production through ZIKV employment of RNase L-cytoskeleton interactions during formation of RFs.

**Figure 7.**
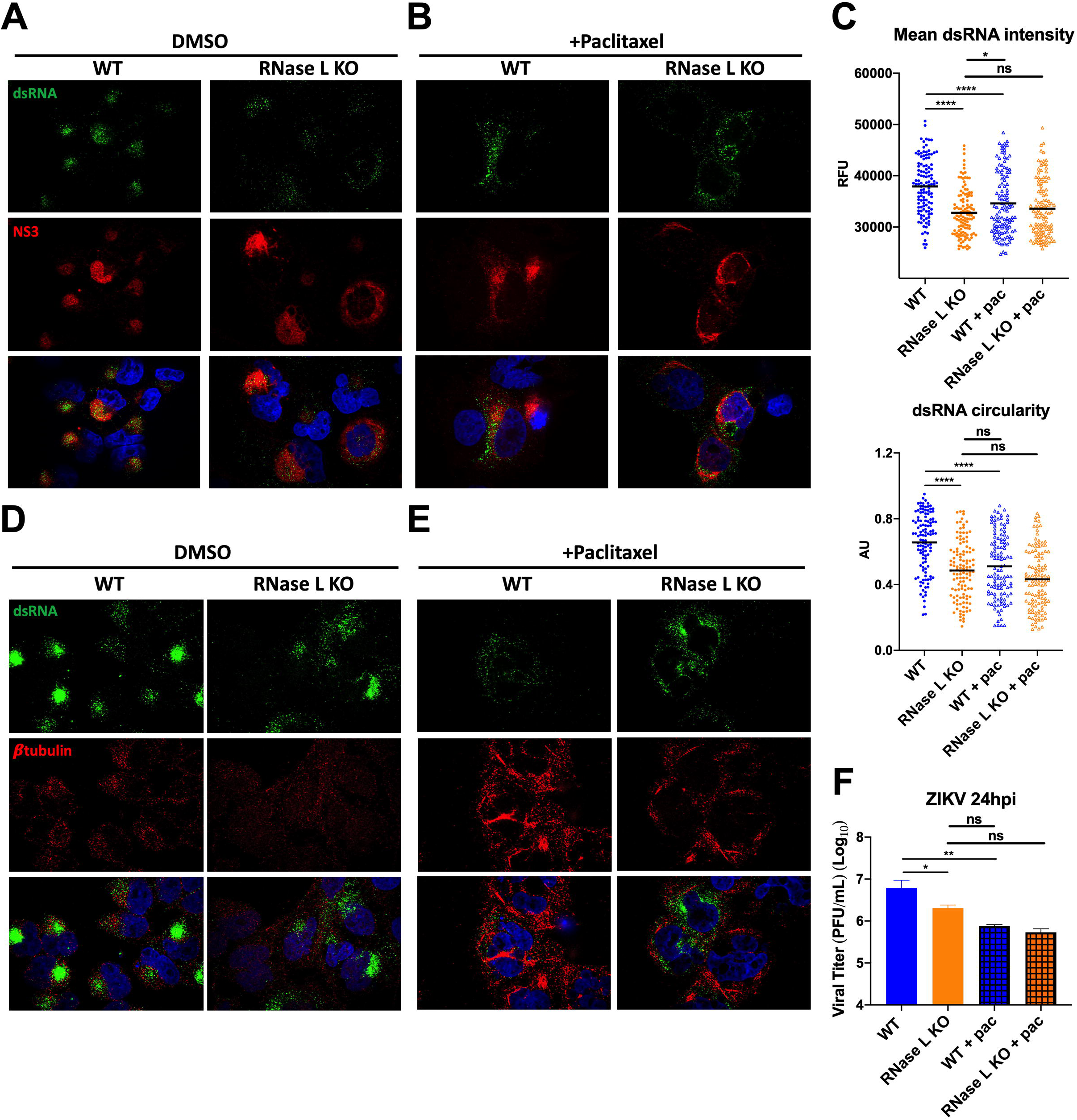
Effects of RNase L deletion on ZIKV RFs resemble antiviral effects of microtubule stabilization on ZIKV RFs. A549 WT or RNase L KO cells were infected with ZIKV at an MOI of 1, treated with 12.5μM paclitaxel or DMSO 3h after infection. For IFA staining cells were fixed at 20hpi. Cells treated with (A) DMSO or (B) paclitaxel were stained for dsRNA (green) and ZIKV NS3 (red), with DAPI (blue) nuclei staining, mean dsRNA intensity and circularity was quantified in (C). Cells treated with (D) DMSO or (E) paclitaxel were stained for dsRNA (green) and βtubulin (red), with DAPI (blue) nuclei staining. (F) For infectious virus production, supernatants were harvested at 24hpi for measurement of viral titers by plaque assay, shown as plaque forming units (PFU)/mL virus. Data is representative of at least two independent experiments. Statistical significance was determined by one-way ANOVA, displayed is the mean of three replicates ± SD for replication assays, ns = not significant, *p<0.05, **p<0.01, ****p<0.0001. Imaged at 100X magnification. RFU = relative fluorescence units, AU = arbitrary units.

## Discussion

In this study, we present findings supporting a proviral role of the host RNase L protein, which has otherwise been considered strictly antiviral. RNase L nucleolytic activity is highly effective at restricting replication of a diverse range of DNA and RNA viruses (Silverman, 2007). However, catalytic activation of RNase L by ZIKV does not limit infection, and instead ZIKV titers are paradoxically increased with RNase L expression. In particular, a catalytically inactive form of RNase L improved ZIKV RF assembly and function, thereby boosting virus production. We also provide evidence suggesting that ZIKV repurposes the interaction between RNase L and the cytoskeleton to facilitate rearrangement of the ER for establishment of ZIKV RFs. While cytoskeletal arrangements for RF assembly within ER folds are characteristic of flaviviruses, we found that RNase L expression during DENV and KUNV infection had only minimal effects on RF formation. Furthermore, RNase L was ultimately restrictive of both viruses, operating in its canonical antiviral manner to decrease DENV and KUNV titers through its catalytic activity. Therefore, we propose that recruitment of RNase L for enhanced RF function and virus release is a feature presently observed only during ZIKV infection.

Interestingly, DENV production in WT cells was limited by RNase L, as titers increased significantly in RNase L KO cells (Figure 1C), although we were unable to detect any rRNA degradation in DENV-infected WT cells (Figure 1B). Unlike other viruses we have examined, RNase L antiviral effects on DENV were not dependent on OAS3, as viral titers in WT and OAS3 KO cells were similar (Figure 3D). Moreover, RNase L deletion enhanced DENV genome expression (Figure 4B&E), which correlates with increased titers in RNase L KO cells, and suggests that RNase L is activated during DENV infection. Indeed, our previous study indicated RNase L activation and degradation of DENV RNA during infection (Whelan et al., 2019). While rRNA degradation is a hallmark of RNase L nucleolytic activity and therefore a commonly used marker for RNase L activation, it is possible that host rRNA effectively avoids activated RNase L while viral genome is primarily targeted during DENV infection in particular (Li et al., 1998, Rath et al., 2015, Donovan et al., 2017). Though these results present an interesting possibility, there is currently no knowledge of RNase L catalytic function independent of OAS-mediated activation, thus further investigation is required for elucidation of the mechanism behind RNase L restriction of DENV.

Use of pIC to activate RNase L prior to ZIKV generation of protective RFs (Figure 5) was primarily implemented as added measurement confirming RNase L catalytic inactivity during ZIKV infection of OAS3 KO cells, in addition to rRNA degradation (Figure 1B) and 2-5A generation (Figure 3A) assays. OAS3 KO cells therefore provided an elegant tool for individually assaying RNase L catalytic and non-catalytic functions during ZIKV infection (Figures 3&4).

To validate our results in A549 cells, we used HeLa M cells stably expressing RNase L WT, an RNase L catalytic mutant R667A, or an empty vector. Since HeLa M cells do not express endogenous RNase L, the empty vector-expressing cells served as an RNase L-deficient control for these experiments. In addition, HeLa M cells were permissive to flavivirus infection and therefore offered a convenient system for isolation of RNase L catalytic and non-catalytic functions during infection. However, one caveat we encountered was the variance in RNase L WT and R667A protein levels, which occurred when overexpressing these constructs in several cell types. RNase L activation induces apoptosis to limit viral infection, therefore it was important to eliminate the possibility of spontaneous activation of RNase L and subsequent induction of apoptosis in cells stably expressing RNase L WT, which would reduce cell quantity. We verified that RNase L was not activated in uninfected HeLa M cells expressing RNase L WT, as we did not detect any rRNA degradation in mock infected cells (Figure 6B, 3^rd^ column). Moreover, levels of RNase L WT in HeLa M cells were sufficient for rRNA degradation (Figure 6B) and slightly reduced KUNV production (Figure 6E), indicating antiviral RNase L activity. Since we were interested in confirming a non-catalytic function of RNase L, this system was effective for corroborating results in A549 cells, in addition to providing a second cell line in which inactive RNase L enhanced ZIKV production.

After establishing that latent RNase L facilitated ZIKV RF formation, we next sought to link RNase L proviral effects on ZIKV infection with RNase L-cytoskeleton interactions. Interest in cytoskeleton involvement stemmed from an earlier study showing inactive RNase L interacted with the cytoskeletal protein Filamin A for inhibition of viral entry. That study also determined that this interaction required RNase L latency, and was dissociated upon catalytic activation of RNase L (Malathi et al., 2014), which correlates with our proposed function of inactive RNase L during ZIKV infection. Another study described the cytoskeleton-containing cellular fraction as also comprising a form of RNase L that was unresponsive to catalytic activation by 2-5A (Tnani et al., 1998). Proteomics analysis in a human cell line identified IQGAP1 (an IQ [isoleucineglutamine] motif containing GTPase activating protein 1), a mediator of actin cytoskeleton reorganization, as an RNase L binding partner (Sato et al., 2010), and another report revealed 62% of RNase L interactions in mouse spleens were with cytoskeleton and motor proteins (Gupta and Rath, 2014). Additionally, RNase L interacts with a regulator of tight junctions, ligand of nump protein X (LNX) (Ezelle et al., 2016), although none of these findings were in the context of virus infection.

While RNase L function during ZIKV infection is the first suggestion of a proviral inactive RNase L role, the aforementioned latent RNase L interaction with Filamin A indicated that RNase L can actively remodel cytoskeleton during viral infection, which is consistent with reports of virus manipulation of cytoskeletal components to facilitate virus replication cycle events (Foo and Chee, 2015, Walsh and Naghavi, 2018). It is known that ZIKV induces extensive cytoskeletal remodeling during infection to establish RFs within invaginations of the ER (Cortese et al., 2017). Since we found that RNase L deletion altered the localization of RF constituents, including ZIKV dsRNA and NS3, ER, and βtubulin, to less concentrated areas that generated less virus, we hypothesized that this was due to a loss of cytoskeletal remodeling without RNase L. Although we attempted to examine alterations in latent RNase L localization upon ZIKV infection, this proved difficult without an effective antibody for visualizing RNase L by IFA. Furthermore, attempts at IFA detection of tagged RNase L localization during infection were inconclusive with unequal overexpression levels of RNase L WT and R667A. We instead used the microtubule stabilizing drug paclitaxel to recapitulate the ZIKV RF phenotype of RNase L KO cells. Paclitaxel-mediated loss of microtubule flexibility disturbed ZIKV RF assembly and function and consequently reduced virus production, as was observed in RNase L KO cells (Figure 7). These results supported ZIKV employment of RNase L’s association with the cytoskeletal to generate RFs (Figure 7). Distinctions in host protein composition of ZIKV RFs compared to that of other flaviviruses have been described (Cortese et al., 2017). Since RNase L improves production of ZIKV but not DENV or KUNV, it is possible that RNase L is another ZIKV RF component not shared among all flaviviruses. Inactive RNase L participation in ZIKV RF formation also implies lower availability of activated RNase L proximal to ZIKV RFs, which offers explanation for ZIKV protection from antiviral effects of RNase L, while DENV and KUNV are sensitive to RNase L activity.

Despite clear alterations in ZIKV RF shape and function without RNase L facilitation of cytoskeleton remodeling, ZIKV RFs still form and function to a degree in the absence of RNase L. This suggests redundancy in host protein manipulation for cellular remodeling by ZIKV, and is supported by the existence of RFs even after RNase L is activated and cleaving ZIKV genome outside RF sites. Additionally, the dissemination of ZIKV RFs in RNase L KO cells occurred in a majority of infected cells, but not in every infected cell, indicating that other host proteins may compensate for the loss of RNase L. Further studies examining RNase L interaction with specific ZIKV proteins or genome are required to identify if RNase L is directly involved in assembly of ZIKV RFs.

## Acknowledgements

We thank Erick R. Perez and Courtney Comar for their help with KUNV and image quantification experiments, respectively, and Dr. Andrea Stout and the Cell and Developmental Biology Microscopy Core at the University of Pennsylvania for confocal microscopy training. This work was supported by NIH grants R21NS100182 (to S.R.W) and R01AI104887 (to S.R.W. and R.H.S), and R01AI135922 (to R.H.S.). J.N.W. was supported in part by T32NS007180.

## Authors Contributions

J.N.W., R.H.S., and S.R.W. designed the experiments. J.N.W., J.H., D.M.R. and B.D. performed the experiments. J.N.W. wrote the manuscript. J.N.W, R.H.S., and S.R.W. edited the manuscript. S.R.W supervised all research.

## Methods

### Cells and viruses

A549 cells were grown in Roswell Park Memorial Institute media 1640 (Gibco 11875) supplemented with L-Glutamine and 10% (vol/vol) fetal bovine serum (FBS, Hyclone*^®^*). Vero cells were grown in Dulbecco’s modified eagle’s media (DMEM, Gibco 11965) supplemented with 4.5g/L D-Glucose, 10% FBS, 10mM HEPES, 1mM sodium pyruvate, and 50µg/mL gentamicin (Gibco). A549 OAS1 KO, OAS2 KO, OAS3 KO, and two RNase L KO clones were generated as previously described (Li et al., 2016, Li et al., 2017). HeLa M cells expressing pcDNA3 vector, RNase L WT, or RNase L nuclease dead mutant R667A (Dong et al., 2001) were cultured in DMEM supplemented with 10% FBS. Zika virus 2015 Puerto Rico isolate PRVABC59 (KX377337) and Dengue virus type 2 (DENV2, New Guinea C, bei resources) were obtained from Dr. Scott Hensley, University of Pennsylvania, Philadelphia, and were propagated in Vero cells. West Nile virus Kunjin strain MRM61C was provided by Dr. Sara Cherry, University of Pennsylvania, Philadelphia, originally obtained from Dr. Alexander Khromykh, University of Queensland, Australia (Khromykh and Westaway, 1994), and propagated in BHK-21 cells.

### Virus replication assays

Cells were seeded into 24-well plates and infected the next day at indicated MOI in triplicate. After 1h incubation of cells with virus at 37°C, virus was removed and cells washed three times with PBS before addition of 1mL complete media. At indicated time post-infection, 150μl supernatants were harvested and stored at −80°C until titration, and 150μl complete media replaced in each well so that all time points from one replicate contain supernatant from the same infected well throughout the time course. Data is representative of 2 or more experiments.

### Plaque assays

Virus supernatant from infected cells was serially diluted in DMEM supplemented with 2% FBS and added to Vero cell monolayers in 6-well plates. Plates were incubated for 1 hour at 37°C before overlaying infected monolayers with DMEM containing 3% FBS, 8% NaCO_3_, 10mM HEPES, 1X L-glutamine, 250μg/mL amphotericin B, and 0.7% agarose. Plaques were stained with neutral red after 2 days (SINV) or 4 days (ZIKV, DENV) for 16-18h before counting plaques. Viral titers for each triplicate were calculated as plaque forming units per mL (PFU/mL) of supernatant and as the mean of duplicates. All viral titers are displayed in Log_10_ scale as the mean ± SD.

### rRNA degradation assay

A549 cells were mock infected or infected with indicated virus at an MOI of 1 or 5 for the indicated number of hours. Cells were harvested in RLT buffer (RNeasy Mini Kit, Qiagen) and lysed through a QIAshredder (Qiagen). Total RNA was extracted and analyzed on RNA microchips using an Agilent 2100 BioAnalyzer (Xiang et al., 2003, Zhao et al., 2012).

### Immunofluorescence assays

Cells were seeded onto glass coverslips in 12-well plates and the next day mock infected or infected with ZIKV at an MOI of 1. Cells were fixed with 10% formalin at indicated timepoint post-infection (20hpi except for S1A), permeablized with 0.1% triton X-100 in PBS for 15 minutes, and blocked with bovine serum albumin (BSA) for 1h. Coverslips were incubated in primary then secondary antibody diluted in 1% BSA for 1h at RT, with three PBS washes after each antibody. Mouse anti-dsRNA antibody (J2, Scicons) was diluted at 1:500, rabbit anti-protein disulfide-isomerase (PDI, Cell Signaling C81H6) for labeling ER was used at 1:200, rabbit anti-βtubulin (Cell Signaling 9F3) was used at 1:200, and rabbit anti-ZIKV NS3 (GeneTex 133320) was used at 1:1,500. Goat anti-mouse Alexa fluor*^®^* 488 (Invitrogen A11029) and goat anti-rabbit Alexa fluor*^®^* 594 (Invitrogen A11037) secondary antibodies displayed as green and red stains, respectively, were used at 1:400. Cells were briefly incubated with 4′,6-diamidino-2-phenylindole (DAPI) to stain nuclei, then rinsed with PBS. Coverslips were mounted onto glass slides and sealed using ProLong Gold antifade reagent (Invitrogen P36930), and imaged using a Nikon Ti2E fluorescence widefield microscope at 60X (Figure 5) or 100X magnification (Figure S1), or using a VT-iSIM confocal microscope at 100X magnification (all other IFA images). Images shown are representative of 5 or more fields of view taken per condition, for each of 2 or more independent experiments. All images within individual experiments were batch processed using the same contrast and threshold settings before measurement. IFAs for quantification were imaged on a Nikon Ti2E fluorescence widefield microscope at 20X magnification.

### Fluorescence *in situ* hybridization

A549 cells were seeded onto 8-well chamber slides (EMD Millipore) and the next day infected with ZIKV at an MOI of 1. At 20hpi, cells were fixed with 10% formalin for 30 minutes at RT, dehydrated, rehydrated, treated with protease and stained with fluorescent labels specific to positive-strand ZIKV genome (cat 521511), positive-strand DENV genome (cat 528001), or positive-strand West Nile virus genome (cat 475091) using RNAscope® Fluorescent Multiplex Reagent Kit (Advanced Cell Diagnostics, Inc.). After FISH staining, slides were rinsed with ddH_2_O, rinsed with PBS, then blocked with BSA and stained for PDI as described for IFAs, using goat anti-rabbit Alexa fluor*^®^* 488 (Invitrogen A11005) secondary antibody for green ER staining. RNA was imaged on a VT-iSIM confocal microscope at 100X magnification. Images shown are representative of 5 or more fields of view taken per condition, for each of 2 or more independent experiments. All images within individual experiments were batch processed using the same contrast and threshold settings before measurement. Images for quantification were taken on a Nikon Ti2E fluorescence widefield microscope at 20X magnification.

### Image quantification

IFA and FISH images for quantification were taken on a Nikon Ti2E fluorescence widefield microscope at 20X magnification. Scatter dot plots shown are representative of at least two individual experiments with >100 infected cells measured per sample, from at least five fields taken per condition, with mean displayed as a black bar. For IFA quantification, images were processed for quantification using Imagej/FIJI by setting a threshold to define the regions of interest (ROIs) as individual RFs (labeled by dsRNA or NS3). Mean intensity was calculated by measuring the mean gray value within ROIs (RFs). Circularity was calculated using the formula 4pi(area/perimeter^2), where an RF in the shape of a perfect circle results in a value of 1. Feret’s diameter, also known as the maximum caliper, is the largest distance between two points on the perimeter of the RF. Mean gray value, circularity, area, and Feret’s diameter are standard measures of ROIs in ImageJ/FIJI. For FISH images, ROIs contained all RNA throughout each individual cell. Since viral RNA ROIs did not take on a defined shape, a perimeter around each ROI was drawn manually, and area of the ROI measured. Relative fluorescence units (RFU) for intensity, microns for Feret’s diameter, and microns^2^ for area are displayed on the y-axis. Circularity measurements do not have a unit and are reported as arbitrary units (AU).

### 2-5A fluorescence resonance energy transfer (FRET) assays

A549 cells seeded into 6-well plates were infected the next day with ZIKV at an MOI of 5, or mock infected. After 24h, cells were washed with PBS and scraped off of the plate, centrifuged at 1,000 x *g* for 15 minutes at 4°C. PBS was aspirated, and 200μl NP-40 (1%) buffer pre-heated at 95°C for 2 minutes was added to pellet. Samples were further processed as described (Gusho et al., 2014) and assayed for 2-5A levels by FRET assays as described (Thakur et al., 2005).

### pIC transfection

Cells were seeded in 24-well plates, with coverslips if for immunofluorescence assays (IFA), and the next day transfected ± 250ng/mL pIC using Lipofectamine 2000 (Invitrogen) for 2h prior to infection with ZIKV at indicated MOI. Cells not treated with pIC were transfected with Lipofectamine only. Cells were either fixed for IFAs at 24hpi or supernatant for replication curves was harvested at 2 and 24hpi for downstream analysis.

### Paclitaxel treatment

Paclitaxel (Sigma-Aldrich) was resuspended in dimethylsulfoxide (DMSO). Cells were infected with ZIKV at an MOI of 1 for 3h before paclitaxel or DMSO was added to cell supernatant at a final concentration of 12.5μM, as described in (Cortese et al., 2017). Cells were fixed or supernatant harvested for downstream analysis.

### Western blotting

Whole cell lysate was boiled for 10 minutes at 95°C, loaded onto pre-cast 10% polyacrylamide gels (Bio-Rad), migrated through the gel by SDS-PAGE at 110V, transferred onto PVDF membrane (EMD Millipore) for 3h at 150mA, blocked in 5% milk overnight, incubated in primary and HRP-conjugated secondary antibodies in 5% milk for 1h each incubation, and imaged using chemiluminescent HRP substrate (Thermo Scientific™ SuperSignal™ West Pico). Goat anti-OAS3 N-18 (Santa Cruz) was used at 1:250, mouse anti-human RNase L (Dong and Silverman, 1995) at 1:1000, and anti-GAPDH as a loading control at 1:1000 (Thermo Fisher MA5-15738).

### Statistical analyses

All analyses were performed in GraphPad Prism version 8.2.1. Plaque assay data comparing WT and RNase L KO groups only were analyzed by student’s paired t test. Image quantification data comparing WT and RNase L KO groups only were analyzed by student’s unpaired t test. Plaque assay and image quantification data comparing three or more groups, and 2-5A FRET assay data were analyzed by one-way ANOVA. ns = not significant, *p<0.05, **p<0.01, ***p<0.001, ****p<0.0001.

