## Supplemental figures S1 and S2 for "The host antiviral ribonuclease L protein supports Zika virus replication factory formation to enhance infectious virus production"

**Figure S1. Loss of RNase L causes ZIKV RNA and protein dissemination.** A549 WT cells and two different RNase L clones were infected at an MOI of 1, cells were fixed at 12, 16, 20, or 24hpi for IFA staining for dsRNA (green) and ZIKV NS3 (red), with DAPI (blue) staining of nuclei. Data is representative of at least two independent experiments. Imaged at 100X magnification, scale bar 20μm.

**Figure S2. RNase L has minimal effects on DENV and KUNV dsRNA expression in the absence of OAS3.** A549 WT, RNase L KO, or OAS3 KO cells were infected with DENV or KUNV at an MOI of 1, fixed at 20hpi, and stained by IFA for dsRNA (green) for quantification of dsRNA mean intensity, circularity, and diameter. Data is representative of at least two independent experiments. Statistical significance was determined by one-way ANOVA, ns = not significant, *p<0.05, **p<0.01, ***p<0.001, ****p<0.0001.
